# GPCR-IFP: A Web Server for Analyzing G-Protein Coupled Receptor-Ligand Fingerprint Interaction from Experimental Structures

**DOI:** 10.1101/2022.03.16.484667

**Authors:** Marc Xu, Yi li, Junlin Dong, Shuguang Yuan

## Abstract

G-protein coupled receptors (GPCRs) are interesting targets for a broad range of drugs. The comprehensions of the interaction’s mode between GPCRs and their ligands remains to be resolved. Here, we present GPCR-IFP, a web server which drives to gather experimental information supports a better understanding of ligand-receptor interaction and the chemical space. Despite the vast chemical library used for drug discovery, the structures of ligands for a particular receptor share some common molecular features. Thus, we foraged to collect all experimental structures of GPCRs and analyzed protein-ligand interactions which resulted in important unique patterns for both agonist and antagonist. All those results can be analyzed and visualized in GPCR-IFP, a friendly-used webserver.

## INTRODUCTION

G-protein coupled receptors (GPCRs) are one of the most considerable families of receptors embedded in cell membranes. The particularity of these receptors is well-characterized by their seven transmembrane alpha-helices (TM) connected by intracellular and extracellular loops. The extracellular domain of GPCRs generally entails an orthosteric binding site allowing ligand binding which in turn triggers downstream heterotrimeric G-protein [1] coupling to the receptor intracellular domain. Despite all GPCRs share a consensus serpentine topology, they involve in a broad panel of pharmacological properties and roles in various physiological processes.

GPCRs are targeted extensively by various pharmaceutical drugs for a vast number of diseases. The Food and Drug Administration (FDA) has reported that about 30 – 40% of approved drugs targeted directly at GPCRs [2–5], making GPCRs a popular and prominent target for drug discovery. GPCRs respond to the ligand binding and transduce the extracellular signals into intracellular responses [6]. In addition, some poly-pharmacological drugs unexpectedly act on GPCRs as secondary targets, provoking or increasing undesirable side effects [7]. The number of GPCR currently reached more than 800 receptors [8], which have been classified into five major classes (glutamate, rhodopsin, adhesion, frizzled, and secretin; GRAFS). This was proposed by Fredriksson [9]. Among all GPCR families, the class A (or rhodopsin-like) receptors are the best well-studied group in both structures and numbers. Class A GPCRs are also in the spotlight of the server in this work.

The Protein Data Bank (PDB) [10] database confers the resource of protein structures, of which more than 77% coupled with small molecules. The binding mode between protein and its ligand requires a specific arrangement of atomic forces via specific conformations in the binding pocket. Through that wealth of data, we can gain insights into how ligands interact with their target proteins as well as the binding site features. On this basis, we intend to design a server capable of analyzing, visualizing, and comparing interactions receptors with different structures and ligands. To achieve this goal, we focus on class A GPCRs which are the most relevant therapeutic targets, of which drug development can be agonists or antagonists. For present and future studies, the mechanism of agonist and antagonist molecules interacting with the orthosteric pocket of GPCRs is an efficient benchmark for designing specifically functional drug molecules.

We propose to manage the resolved receptor structures to develop a tool that is capable of analyzing the protein-ligand interactions. The experimental structures substantially seek out relevant key residues that involved in protein-ligand interactions with complementary computational methods suggesting putative binding mode to explain the ligand activity mechanism. Consequently, computational predictions become challenging due to the fact that particular GPCRs have low sequence identity with available GPCR structures [11], and the docking results for estimating receptor and ligand affinity partially include false-positive hits.

In this work, we are focused primarily on compounds that bind family A GPCRs. For this purpose, we established an online server, namely GPCR-IFP (http://cadd.siat.ac.cn/gpcr-ifp/). The server establishes a potential pharmacological profile of drugs. The advantage of the server is to provide visual access to specific (agonist or antagonist bound structure) or unique (exclusively one complex structure) interactions fingerprint between receptor and ligand. Furthermore, different receptor structures have been faithfully aligned. Users can study the conformation changes of the protein pockets.

## RESULTS AND DISCUSSION

The GPCR-IFP server outlines a dynamic representation of protein-ligand fingerprint interaction from experimental structures. According to the accurate structure from experimental outcomes and multiple structures involved for various receptors, the server can measure interaction and furnish an overview of the ligand-receptor relationships. From this perspective, users can readily submit a request easily. The results are structured in two pages. The first page provides a global visualization of the request receptor, containing every structure whether antagonist- or agonist-bound structures, whereas the second page is rather a more detailed illustration of one structure.

The requirement for GPCR-IFP job inquiry is the receptor name or the PDB code. The PDB code found in the server database is related to the receptor, and the suggestion gives all structures related to the receptor name. Pre-calculation are prior executed and the results content will be rapidly displayed to users via related webpage.

### Input and Output and GPCR-IFP

#### Input

A receptor name or a PDB code should be entered necessarily as input. If the receptor exists in the server database, the homepage (Figure 1) will redirect to the results page.

**Figure 1:**
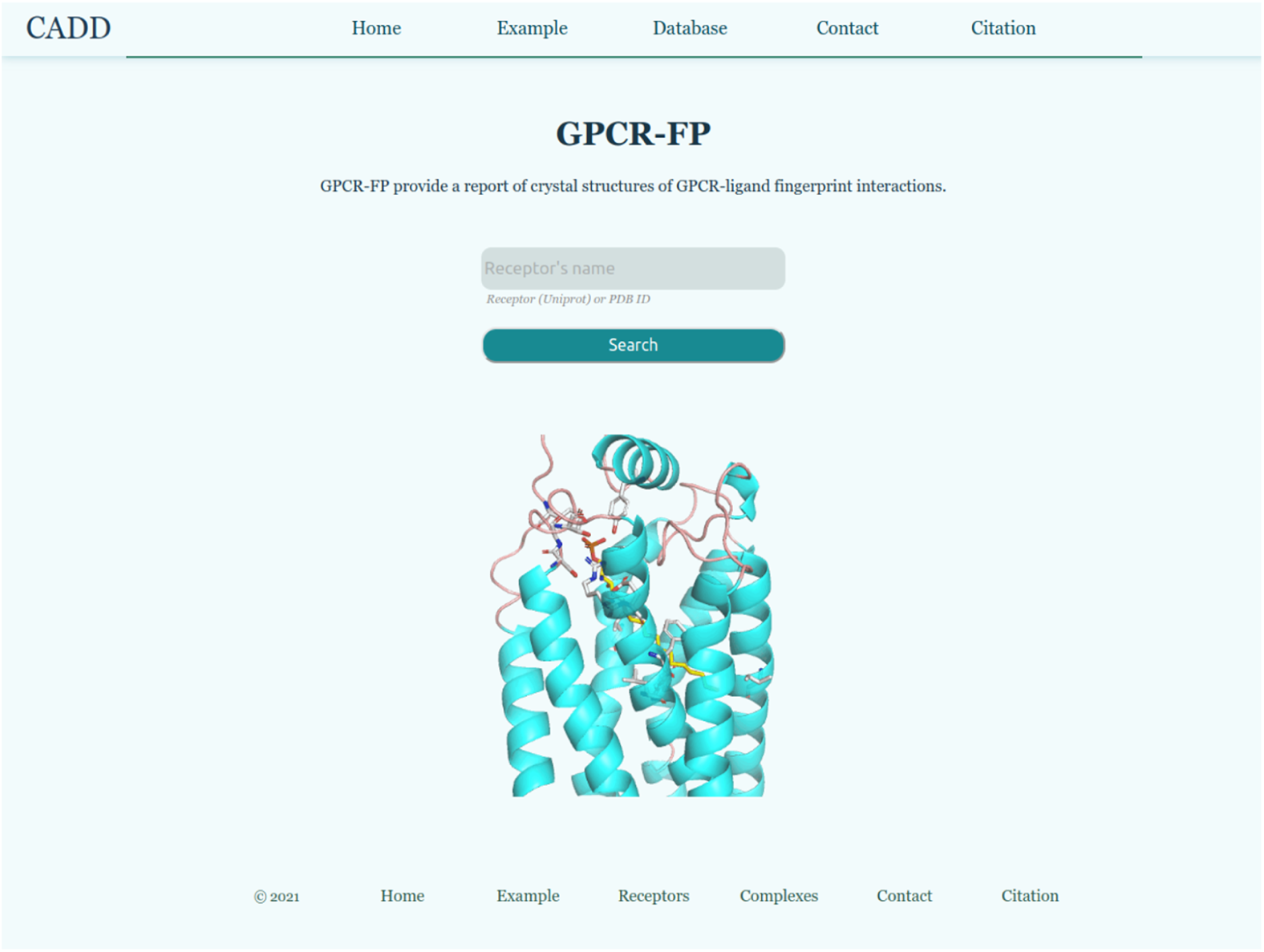
The server homepage.

#### Output

The result contains two pages, namely the global receptor page and the specific structure page.

The jumping page returns the visualization of the cluster of structures for one specific receptor. The page includes agonist and antagonist-target structures, which are displayed in separate windows, and likewise contains the link to reach one particular structure (specific structure with own PDB code).

The global receptor page (Figure 2 left) is subdivided into three parts. The top part enumerates all standard structures of the receptor. The middle part provides the 3D interaction information. The bottom part provides a 2D interaction information map displayed into a radar chart that can download easily. Notably, the viewer of agonist ligand-bound structure and antagonist ligand-bound structure are displayed separately.

**Figure 2:**
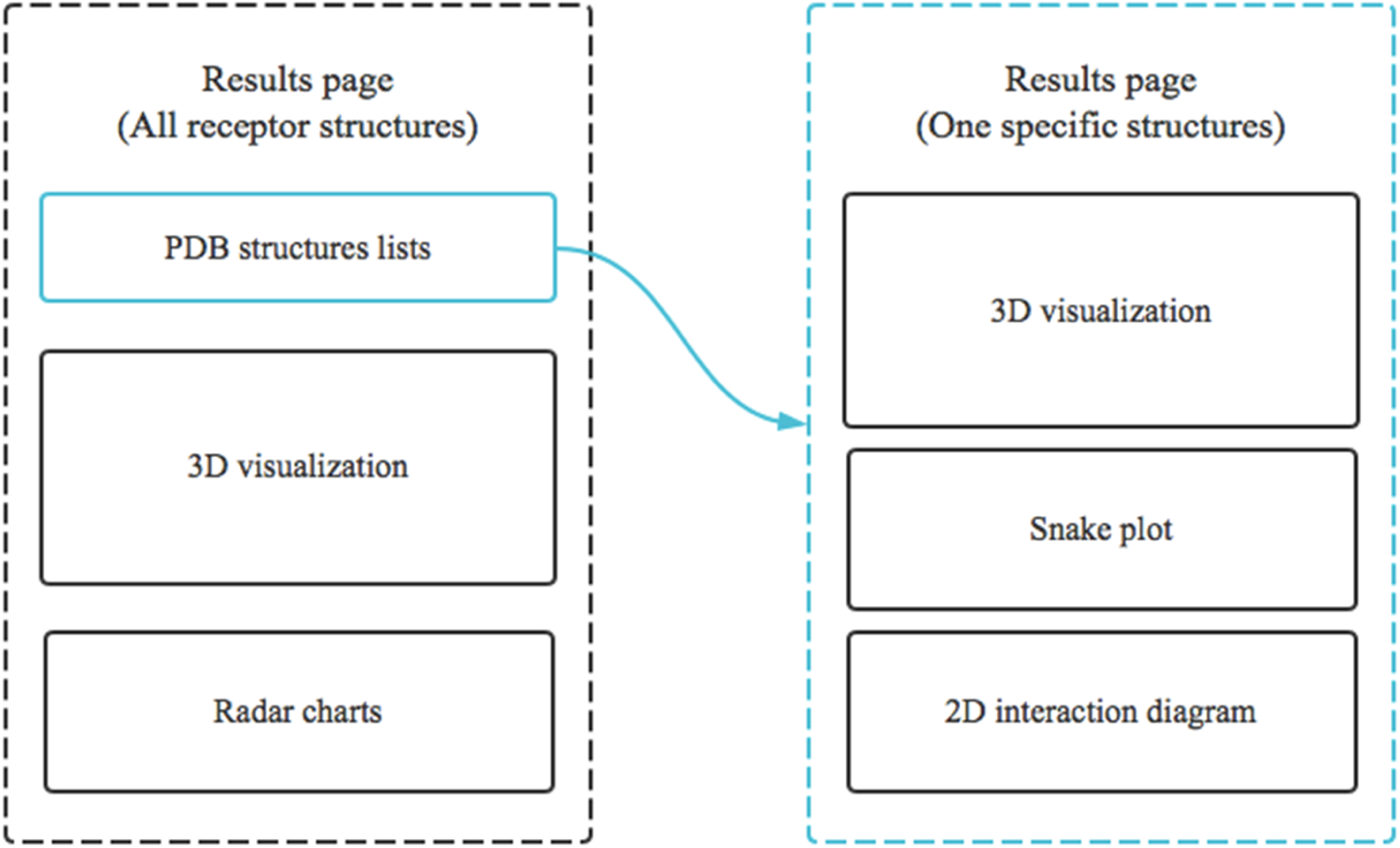
The layout of the results for a given receptor. The left side is the global vision of all structures of a given receptor in 3D visualization and the radar charts of ligand-receptor interaction. The radar chart is the sum of interactions in the complex. On the global results page, structures are represented in cartoons and uniquely colored. The structures with agonist ligand (no representation) are in the left window, while those with antagonist ligand (no representation also) are in the right window. The left side is the results for the specific receptor (recognized by their PDB code) with 3D interaction visualization, the receptor snake plot, and the 2D interaction diagram.

The specific structure page (Figure 2 right) is subdivided into two parts. The top part is the 3D interaction information display. The bottom part incorporates the two plots: the snake plot and the 2D interaction diagram that could directly be downloaded easily in the SVG format. Some complex structures do not enclose a 2D interaction diagram because the Poseview program could not identify the ligand structure correctly.

The functionalities of 3D interaction visualization are listed below:

- Customizing receptors for visualization (global receptor page). There are a couple of selection boxes for all structures of the submitted receptor and two fixed selection boxes to show or not cartoon or sticks representation, respectively, for all structures. The default representation is a cartoon, and users can change the structure in sticks representation. Representations of receptor structures from agonist ligand-receptor complexes and antagonist ligand-receptor complexes are displayed indubitably into different windows. In addition, only receptor structures with the same type of ligand are well-aligned, and corresponding ligands are also represented. To determinate each structure, the color name is indicated after the representation boxes with an asterisk to pinpoint which structure appears in the window.
- Customizing the interest interactions of the complex (specific structure page). This function allows users to select which interaction information will be displayed. All interactions representation is exhibited initially by default. There are nine specific selection boxes for displaying interactions, including eight boxes for specific interaction representation, as well as an option to display all of the interactions or not.
- Customizing the complex structure representations (specific structure page). There is a faculty to visualize a receptor and its ligand in different representations (four representations for receptor and three representations for ligand). The default representation of structures is ligand in sticks and receptor in cartoons. The interacting residues with the ligand are represented even in stick representation that can eventually be hidden.

### Example of application

We selected 5-hydroxytryptamine receptor 2B, 5-HT_2B_ (also known as serotonin receptor) as an example. 5-HT_2B_ is implicated mainly in the cardiovascular system [12]. It regulates many cardiovascular functions such as contraction of blood vessels and shape changes in platelets [13, 14]. Both agonists bound structures and antagonists bound structures were resolved for 5-HT_2B_. The page layout of results is shown in Figure 2. All structures of 5-HT_2B_ are colored distinctively (Figure 3). Structures with the same type of ligands are displayed seamlessly into the same windows spotted by an asterisk and aligned structurally according to their transmembrane (TM) domains. In this specific case, for the agonist-bound structures including 5TVN, 5TUD, 4NC3 and 4IB4 structures have been lined up on the 6DRY. Similarly, 6DS0 and 6DRX, both of which are antagonist-bound structures, are aligned on the 6DRZ. The pairwise distance values showed that structural conformation is very closed between different structures of the 5-HT_2B_ receptor. These structures are not completely transiting on the fully active state, the TM6 structure is an explicit characterization of the receptor state [15] identified between 5TUD (the only active state structure) and all other intermediate state receptors with a slight inward on the cytoplasmic side of TM6 in the active state structure, but no intercalation on the extracellular side.

**Figure 3:**
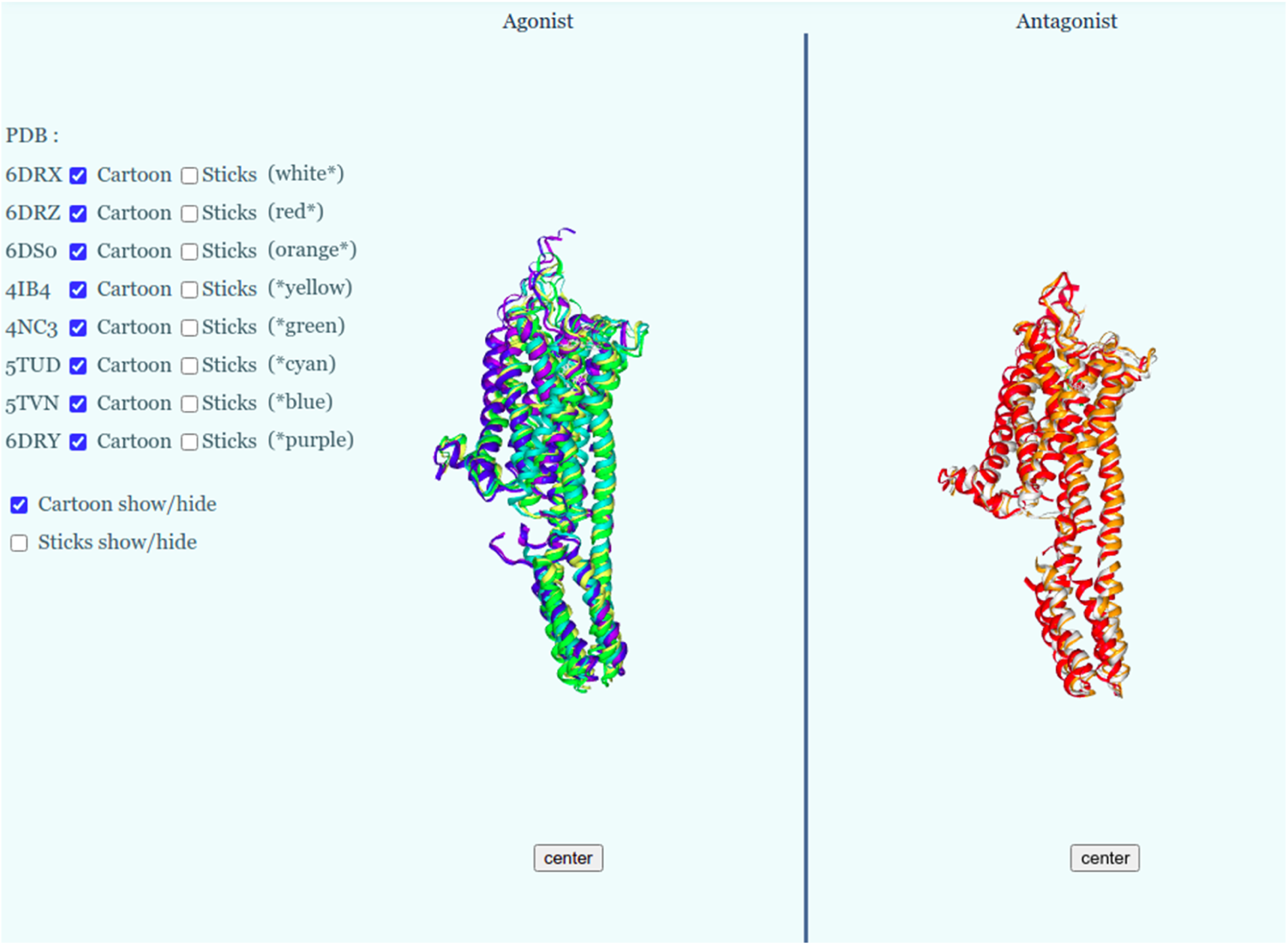
The visualization of structures in three-dimension. The structures are represented in cartoons and colored with a unique color. The structures with agonist ligand (no representation) are in the left window, while those with antagonist ligand (not represented also) are in the right window.

The noncovalent binding mode of each complex is calculated. The sum of interactions is shown in the radar chart (Figure 4). In this plot, all residues involved in the interactions are displayed in the “all” (all interactions) radar chart. Each specific interaction is present in a separate radar chart. The plot shares only the interacting residues and not the count of interaction. In other words, the residue repeatedly interacted with the ligand, the “all” radar chart will consider only once. As an example, the identical residues interacting with two hydrogen bonds upon the ligand binding are weighted evidently as one hydrogen bond. In the case of 5-HT_2B_, the hydrophobic interactions contribute significantly to the ligand-receptor interaction, followed by the hydrogen bonds. The general residues implicated in hydrophobic interactions include V136 and F340, which are found broadly in both agonist and antagonist. Some unique residues are only found in specific ligand types such as L132, F341, Q359, and V366 were mostly in agonist-bound structures. In contrast, L209 and A225 were predominantly in antagonist-bound structures. Other residues in the radar chart have weak frequency suggest that those specific contacts rarely happen. We thus perceive that the number of residues involved in hydrophobic interactions is two-fold higher for agonist ligands-bound structures than antagonist ligands-bound structures for the case of 5-HT_2B_. Hydrogen bonds are slightly more abundant for agonist ligands-bound structures with repetitive contributions of Q135, T140, and L209. In addition, water bridges, an extensive hydrogen bond network, which is linked by water molecules, contributes only to a few structures. Finally, the salt bridges are only found in one structure binding to an antagonist ligand. The binding mode either agonist ligand or antagonist ligand comprised predominantly hydrophobic interactions, supporting ligands are positioned in a deeper hydrophobic cavity to bind with the protein pocket.

**Figure 4:**
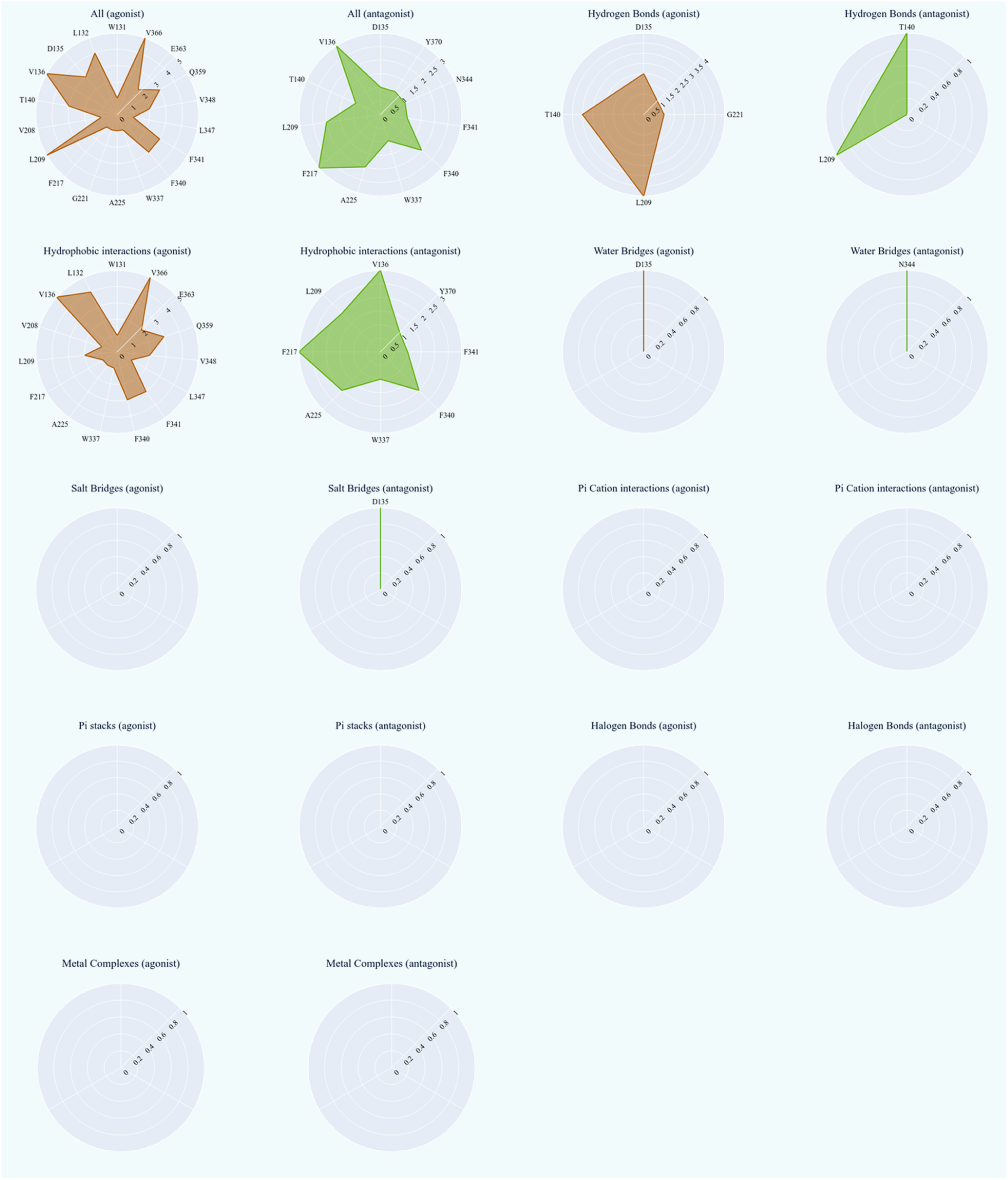
Radar chart summed interactions (all) including hydrophobic interactions, water bridges, salt bridges, hydrogen bonds, pi-stacks, pi-cation interactions, halogen bonds, and metal complexes. These interactions are estimated with the PLIP program. Interaction with agonist ligand (red) and antagonist ligand (green) are positioned next to each other.

After considering interactions for each receptor-pair, molecule ergotamine is found presented in multiple structures (4IB4, 4CN3, 5TUD); both lysergide and the methylergonovine are present in one structure (5TVN and 6DRY, respectively); lisuride is found in two structures (6DRX, 6DRZ) and Ly266097 hydrochloride is in complex with one structure (6DS0). The initial three ligands are agonist ligands, while the last two are antagonist ligands against 5-HT_2B_. The (agonist and antagonist) ligands are formed essentially of one carboxamide or diethylurea, ergoline ring, and their specific terminal moiety (Figure 5). Despite the structural similarity, the three agonist ligands contribute to different therapeutic effects and have been applied to medical treatment distinctively. It prompts us to consider that agonist ligands could activate different pathways inducing various final effects. However, more experimental studies should be executed to validate this assumption for 5-HT_2B_. Although ergotamine is a soluble ligand over others, it exhibits more hydrophobic interactions than other interactions (see in Figure 3). 5-HT_2B_ comprises an extended binding pocket revealing specific ligand binding [16], of which residue M218 has been considered to serve for ligand selectivity [17]. Regardless of the ergotamine moiety, ergoline rings of all ligands are similarly oriented: the indole nitrogen is positioned towards the T140 and A225 permitting the receptor activation [16]. In addition, the three experimental structures of the ergotamine–5-HT_2B_ complex have shown ligand stabilization with different unique interacting residues in their structures: V208 in 4IB4; F217 in 4CN3; L347 in 5TUD, without a particular role in the binding mode. In additional, the residues above-mentioned have covered up the hydrophobic interactions, giving potential binding site features.

**Figure 5:**
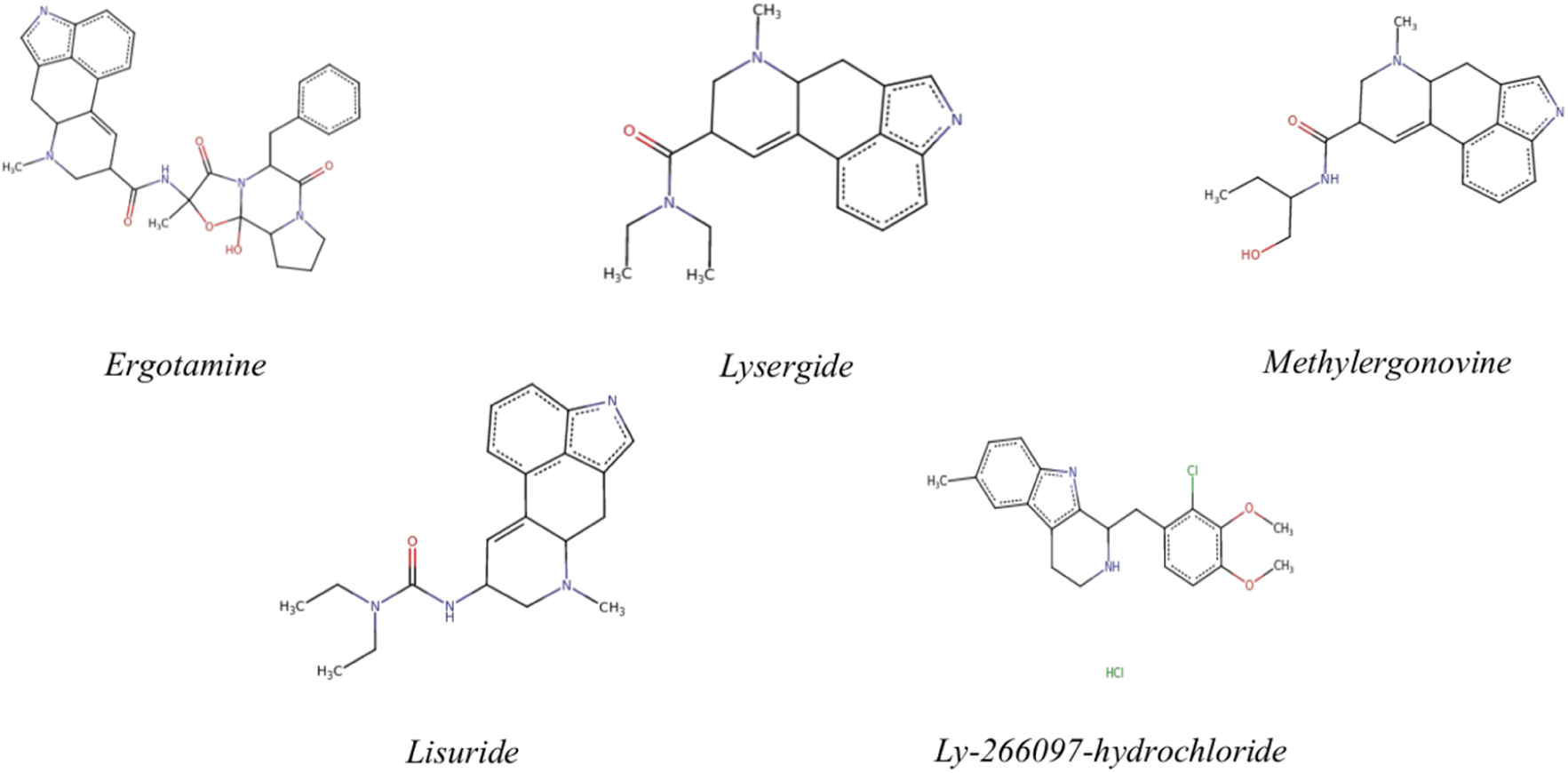
Chemical structure of ergotamine, lysergide, methylergonovine, lisuride and Ly 266097 hydrochloride.

The specific receptor results page also displays a 3D representation (Figure 6) with ligand structure and interaction representations implemented. There is a deeper understanding of the ligand in the binding pocket. The snake plot and the 2D interaction diagram (Figure 7) are suitable implements to describe the evidence. The snake plot (Figure 7, top) stands for the wavy structure of GPCR which residues of the TM domains are represented in circle shape while residues in the loop are illustrated routinely by lines. Residues involved in the stability of the ligand are colored in red. The 2D diagram (Figure 7, bottom) displays the ligand structure in the center, and the residue interactions surface is described by the residue label and the interaction type. This diagram depicts hydrophobic interactions, hydrogen bonds, and pi interactions (pi-stacks and pi-cation interactions). Despite the methods of PLIP [18] and Poseview [19] for calculating interactions being unique in their algorithms and thresholds, the result of the significant interactions remains identical.

**Figure 6:**
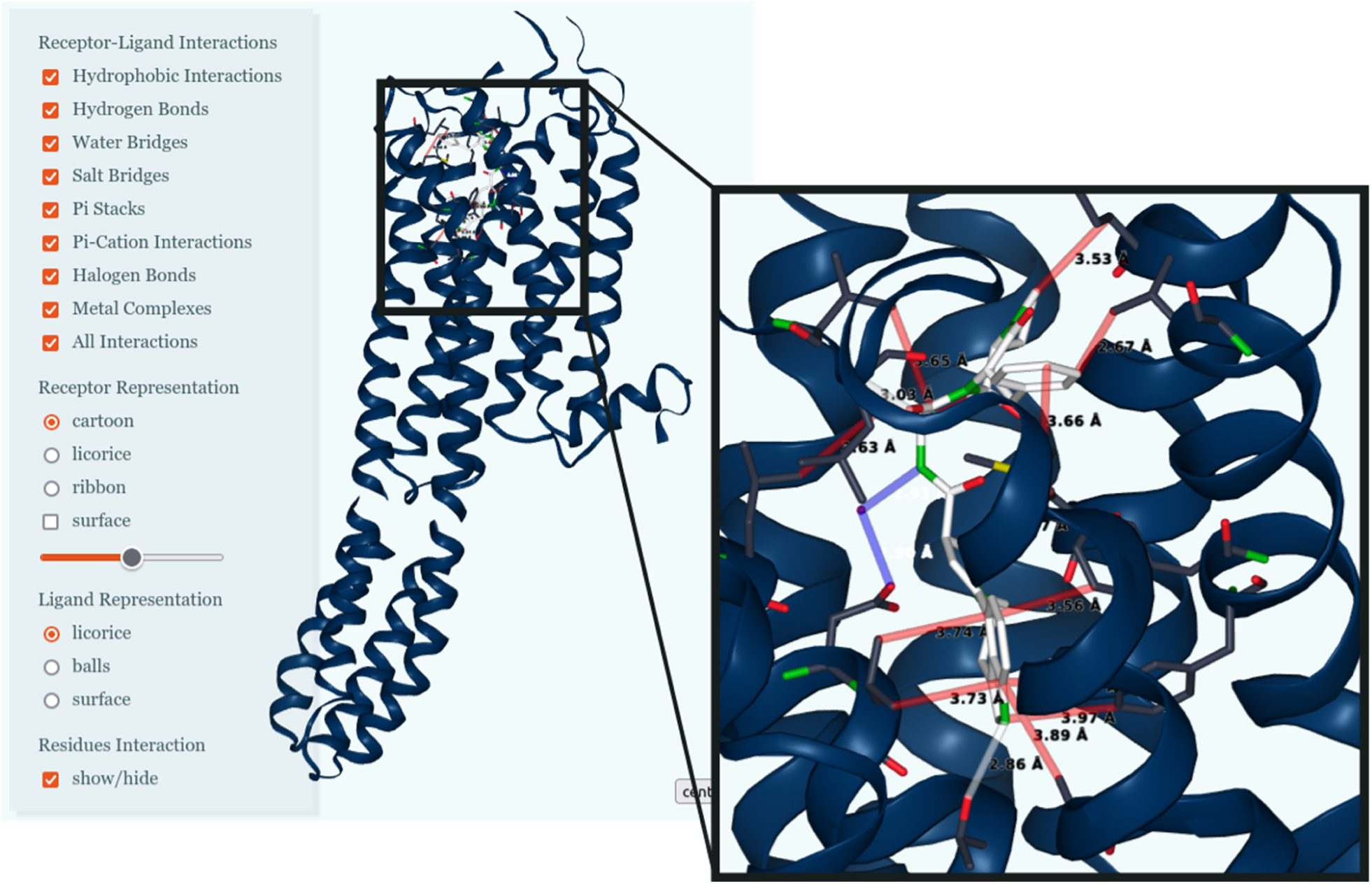
The 3D IFP of the 5-HT_2B_ receptor and ergotamine (PDB code: 4IB4) interaction in GPCR-IFP server. Ergotamine is represented as white sticks, whereas 5-HT2B is represented as blue cartoon. Interactions are indicated by line.

**Figure 7:**
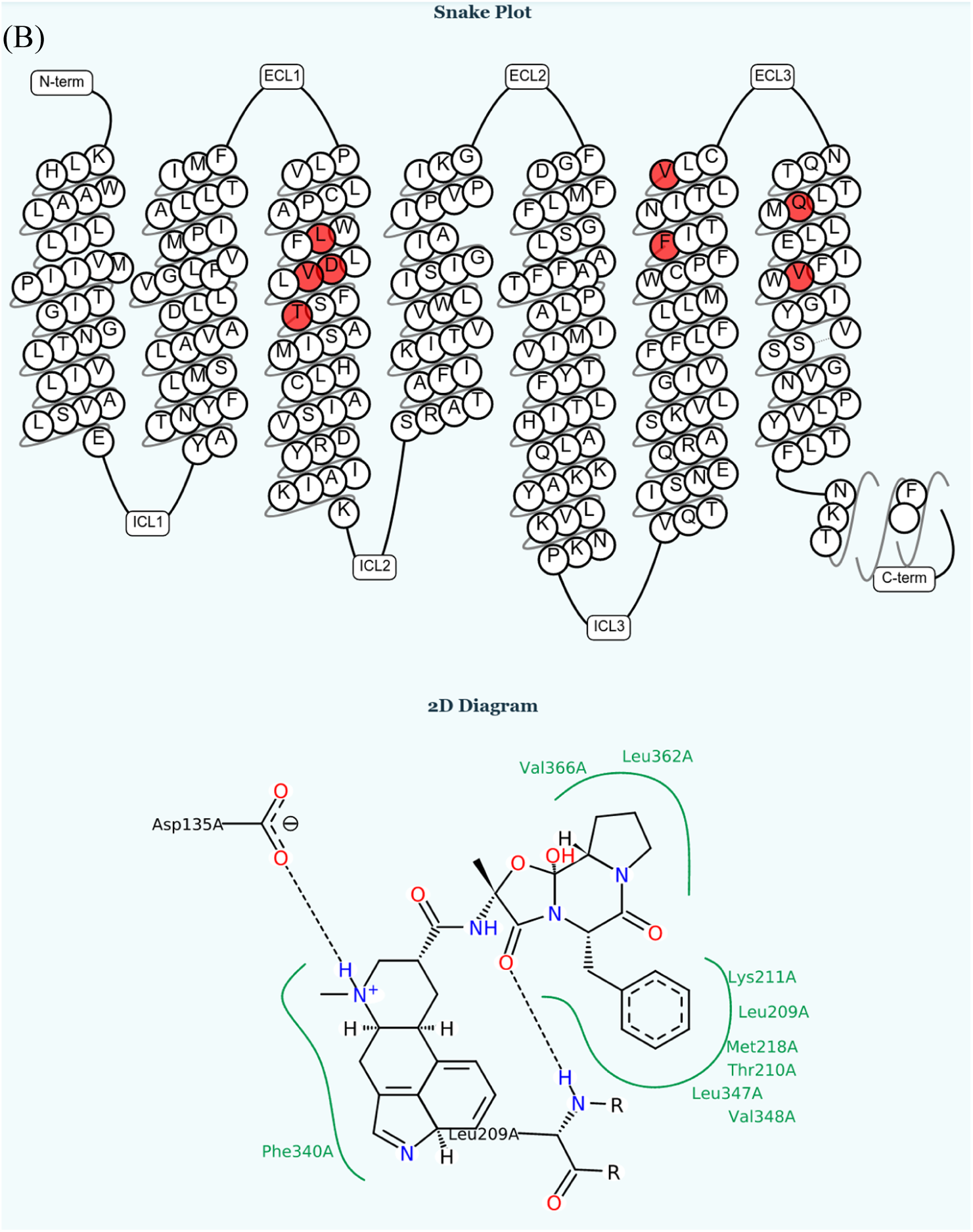
The 2D representation of GPCR-IFP for the interaction between 5-HT_2B_ receptor and ergotamine (PDB code: 4IB4). Top: snake plot of the receptor, the transmembrane residues are represented, and interacting transmembrane residues are colored in red. Bottom: the 2D interaction diagram displays residues interacting with the ligand structure. The green lines and green residues are hydrophobic interactions; and the black dot lines are hydrogen bonds.

Some structures of agonist or antagonist ligand-receptor complexes are in an intermediate state such as structure 4IB4. The deeper insight into different states of class A GPCRs has crossed the milestone thanks to the determination of the first ligand-free experimental structure of opsin receptor [20]. Thereafter, the intermediate state structures combined with spectroscopy and molecular dynamic simulations studies have been provided evidence of the link between the fully inactive and the fully active state conformations [21]. In other words, the receptor activated by the ligand undergoes a conformational change, particularly the binding site at the cytoplasmic regions in order to transmit the signal to the G protein followed by signaling cascade. The knowledge regarding the intermediate state still resided as a vague but unquestionable vision. It was discovered that this state possessed properties similar to the active state, it was manifested in the activation pathways and modulated the G protein [22] which has previously been assigned exclusively for the active state. The intermediate state leads either to the active state or continues to be regulated. Few grounds support this second instance, but we assume that the conditions to fulfill this situation would be allosteric modulation. Previous studies have shown that the absence of the ligand could also undergo transition inactive-to-active state of GPCRs and activate the downstream system but less frequently [8, 23], which suggests the orthosteric ligand-binding increases activity of these receptors and alternative ways to engage the receptor in activate state. The triggered factor of receptor activity could be the membrane lipid (as cholesterol in the experimental structure) interacting on the allosteric binding site buried in the intramembrane regions and regulating the GPCR dynamics and functions [24]. In addition, the hydrophilic ligand could interact and stabilize more straightforward with the extracellular surface, suggesting another potential allosteric binding site [25]. These binding sites confer the allosteric modulators of GPCRs which provides new opportunities for the drug discovery along with novel features on the drug action or pharmacological profile.

The success of a fully active state crystallization is more complicated which needs to stabilize the receptor by an intracellular binding partner, either a G protein or a G protein mimetic that binds on the intracellular regions of an activated receptor [8]. For instance, some structures engineered with flavodoxin could increase broadly the yield and the stability of the protein, which are both significant important for the quality of the final experimental structure [26].

We have established a threshold resolution limit at 3.5 Å and 3.9 Å for x-ray diffraction method and electron microscopy method, respectively, value to select an accurate structure, because a higher resolution structural yields more reliable result. In contrast, unsatisfactorily resolved structures have suspicious side chain conformation and induce uncertainty for the protein-ligand interactions. A broader diversity of ligand would thus enable further basic understanding and therapeutic intervention. Nonetheless, the knowledge and number of experimental structures of GPCRs are being increased continuously which facilitate the understanding of molecular mechanism for agonist and antagonist bindings.

## CONCLUSION

We have established a threshold resolution (3.5 angstroms and 3.9 angstroms for x-ray diffraction method and electron microscopy metthod, respectively) value to select an accurate structure, because a higher resolution structural yields more reliable result. In contrast, unsatisfactorily resolved structures have suspicious side chain conformation and induce uncertainty for the protein-ligand interactions. A broader diversity of ligand would thus enable further basic understanding and therapeutic intervention. Nonetheless, the knowledge and number of experimental structures of GPCRs are being increased continuously which facilitate the understanding of molecular mechanism for agonist and antagonist bindings. The breakthroughs of novel drugs against GPCRs and new insights into these therapeutic targets over the past several years have elucidated many puzzles in this area. The combination of experimental techniques and computational tools enhances GPCR new drug discovery. Although resolving GPCR experimental structures remains a great challenge, many breakthroughs have made in recent years. In addition, computational methods are being investigated to explore putative protein structures that are hard to stabilize. The present fashion of putative three-dimensional structures discovery of GPCRs is homology modeling, ab initio techniques [27] as well as artificial intelligence such as AlphaFold2 developed by Google’s team [28]. Among those resolved structures, GPCRs share commonly three different states: the active state, the intermediate state as well as the inactive state. The active state predominantly triggers downstream signaling.

The protein-ligand interactions fingerprint calculated from the experimental structures yields many valuable information to guide GPCR drug discovery. It not only tells which amino acids involved in drug molecule interactions, but also convey the information that what is the unique features for agonist binding or antagonist binding against a specific GPCR. To achieve this goal, the GPCR-IFP was built. In GPCR-IFP server, it focuses on all experimental structures of class A rhodopsin-like receptors. GPCR-IFP generates characteristic feature of the interaction fingerprint between GPCRs and their ligands in both 3D representation and 2D plots. With those helpful information, one can design a drug candidate with specific functions. GPCR-IFP provides a useful insight into modern GPCR drug discovery.

## AUTHORS CONTRIBUTIONS

MX and YL performed the studies and the main analysis. MX and YL wrote the first draft of the manuscript and improved by YS and DJ. MX prepared all figures which were later.

## MATERIALS AND METHODS

### Overall structural and technology of GPCR-IFP

Our developed an online sever GPCR-IFP to produce interaction fingerprint between GPCRs and their ligands. The non-bonded interactions in both two-dimension and three-dimension graphics are present. GPCR-IFP is implemented with the Django web framework under Python version 3.8. The main python packages were Biopython, Biopandas, and Pymol used to parse PDB files.

### Data collections

The data collection is focused exclusively on family A GPCR structures that are well-documented in the GPCRdb [29]. The completeness of the database affords to retrieve 200 experimental structures of GPCRs (GPCRdb; December 20, 2021). Among collected structures, improper structures have been sequentially released from further analysis, such as flawed resolution structures (resolution of x-ray diffraction method and electron microscopy method upper 3.5 angstroms and 3.9 angstroms, respectively), inactive state structures with agonist molecules, protein-bound structures, and unbound molecule structures. We reached 114 structures with their information in the end, including the receptor name, the host species, the preferred chain, the amino acid sequence, the references to the literature, and basic knowledge of the ligand.

### Structure preparation

Small molecules are recorded as heteroatom in structure files such as water molecules and ligands. These small molecules are filtered thoroughly to identify the correct ligand, whereas non-ligand small molecules such as helper molecules (i.e., 1-Oleoyl-R-glycerol, code name: OLC), cofactors, and ions atoms are extracted unexceptionally out of the structure file. The ligand is identified primarily with the GPCRdb web server, and the information is completed and reviewed by RDkit tools [30], IUPHAR/BPS Guide to Pharmacology database [31], and PubChem database [32]. Hence, the ligand table enclosed ligand properties (molecular weight, number of hydrogen donor/acceptor, function), ligand identification (SMILE, IUPAC name, synonyms…), and the related PDB code. The final structure file includes a receptor, a ligand as well as internal water molecules (forming potential water bridges).

After filtering structures files, we prepared receptor structures and ligand structures. The initial structures do not contain any hydrogen atoms and some side chains might be missing, although the resolution is acceptable. For the protein structure preparation, we executed it in Maestro tool (version 2020.3) [33]. The receptor structure is filled with missing side chains residues and H-bond interactions were optimized.

Despite most ligands are localized unsurprisingly in the orthosteric binding site, the binding mode (i.e., residues involving and interaction type) is apparently distinguished by the ligand features. Thus, the complex interactions are calculated efficiently with PLIP [18]. For ligand structures preparation, we have implemented the OpenBabel package [34] to protonate structures. The PLIP interaction results conferred the basis for downstream analysis, notably the water molecules identification. For constructing the fingerprint interactions, the identical receptor structures holding the analogous ligand type were assembled and aligned subsequently, and their number of interactions was afterward summed up. The 3D structures visualization windows are displayed with the NGL Viewer application [35] and the Iview application [36]. The 2D interaction diagram is built promptly by the Poseview [19].

